# Insight into the central and nodal cells transcriptome of the streptophyte green alga *Chara braunii* S276

**DOI:** 10.1101/2023.02.12.528195

**Authors:** Daniel Heß, Anja Holzhausen, Wolfgang R. Hess

## Abstract

*Chara braunii* is an emerging model system for streptophyte terrestrialization and early land plant evolution. In this study, tissue containing nodal cells was prepared under the stereomicroscope and an RNA-seq dataset generated and compared to transcriptome data from whole plantlets. In both samples, transcript coverage was high for genes encoding ribosomal proteins and a homolog of the putative PAX3- and PAX7-binding protein 1. Gene ontology-based functional assignments revealed for the nodal cell sample main upregulated molecular functions related to protein binding, nucleic acid binding and DNA binding. Looking at specific genes, several signalling-related genes and genes encoding sugar-metabolizing enzymes were found to be expressed at a higher level in the nodal cell sample, while photosynthesis-and chloroplast-related genes were expressed at a comparatively lower level. We detected the transcription of 21 different genes encoding DUF4360-containing cysteine-rich proteins. The data contribute to the growing understanding of charophyte developmental genetics by providing a first insight into the transcriptome composition of *Chara* central and nodal cells.

## INTRODUCTION

Originally mainly considered as a model organism for modern plant electrophysiology (reviewed in Beilby, 2016), studies on plant polarized growth and gravity sensing (Braun and Limbach, 2006), the Charophyceae alga *Chara braunii* Gmel. 1826 (Gmelin, 1826) gained substantial interest, more recently as a model organism to study early land plant evolution. The *Chara* model produces both a high number of oospores within a few months, and sports a short annual life cycle compared to other Charophyceae species. Reproduction can occur generatively by means of oospores, but can also proceed vegetatively by fragmentation of thalli (Casanova, 2015; Holzhausen et al., 2022). The sequenced draft genome (Nishiyama et al., 2018) revealed evolutionarily significant, plant-like features as well as a striking secondary complexity. Furthermore, standardised protocols for vegetative and generative cultivation have been established, enabling a more detailed insight through transcriptomic studies (Holzhausen et al., 2022).

Charophyte algae within the higher branching ZCC (Charophyceae, Coleochaetophyceae and Zygnematophyceae) clade are the closest living relatives to recent embryophytes (Vries and Archibald, 2018; Nishiyama et al., 2018). While Zygnematophyceae are the sister group to land plants (Ruhfel et al., 2014; Wickett et al., 2014; Cheng et al., 2019), the earlier branching Charophyceae form the only class of streptophyte algae with tissue-like structures and rhizoids (Bonnot et al., 2019), traits that were probably already possessed by the streptophyte algal ancestor at least in a rudimentary fashion (Fürst-Jansen et al., 2020).

Roughly 500 million years ago a charophyte progenitor colonized land and gave rise to land plants (Martin and Allen, 2018). The extant Charales, such as *Chara braunii*, bear striking similarities to the morphology of land plants, yet display also important differences in cell biology (reviewed in Domozych and Bagdan, 2022). Therefore, analysis of the *Chara braunii* developmental program can shed light on the evolutionary adaptions associated with these morphological features.

The monoecious macroalgae grows from a terminal apical cell, and consists of a central stem with branchlets radiating from axial nodes at regular intervals. Nodal and elongated, multinucleate internodal cells are derived from the apical cell in alternating fashion, separating each whorl by an internodal cell. The nodes contain at their center a pair of small cells, called central cells and additional 6 to 20 cells that surround this pair. All these nodal cells derive from a single nodal initial cell (Kuczewski, 1906; Schubert et al., 2016). However, these nodal cells can also form apical cells *de novo* that can develop into lateral branches (Nishiyama et al., 2018) and therefore suggest a stem cell-like character. Within a whorl, each branchlet is derived from an initial nodal cell. Along its branchlet nodes, the resulting *Chara braunii* bears both male (antheridia) and female (oogonia) gametangia (Nishiyama et al., 2018; Moody, 2020).

Generally, a comprehensive understanding of early molecular innovations towards the dynamic terrestrial environment is still limited (Zhang et al., 2022). Investigating the developmental genetics of emerging charophyte model systems can shed light on the evolutionary adaptions needed for terrestrialization.

Characeae present certain evolutionary novelties of streptophyte algae, leading to three-dimensional growth: apical cell growth, tip growth and division plane rotation (involving the phragmoplast and cell plates). The nodal cells, found at morphologically key positions between the long internodal cells and branchlets along central cells, can still undergo cell divisions. In contrast to internodal cells, this enables them to generate distinct nodal discs and the founder cells for branches (reviewed by Buschmann, 2020). Compared to oogonia and antheridia, this de novo formation of apical cells also enables asexual propagation (Nishiyama et al., 2018). Here, we performed a transcriptome analysis of a *Chara braunii* sample enriched for nodal cells compared to total plantlet material, providing first insight into the gene expression in these cells. Whole transcriptome sequencing of *Chara braunii* central and nodal cells revealed the relatively higher expression of several signalling-related genes and of genes encoding sugar-metabolizing enzymes, while photosynthesis- and chloroplast-related genes were expressed at a comparatively lower level.

## MATERIALS AND METHODS

### Culture of *Chara braunii*, growth conditions and dissection of central and nodal cells

Vegetative cultures of the non-axenic freshwater strain *Chara braunii* S276 (KU-MACC), originating from Lake Kasumigaura (Ibaraki, Japan), maintained at Kobe University (Sato et al., 2014; Kawai et al., 2022) and consecutively propagated at the Universities of Marburg and Freiburg, were used. Plantlets were grown using the protocol of Holzhausen et al. (2022) in double autoclaved culture vessels containing sieved compost (Gardol® Pure Nature, Bauhaus), lime (Gardol® Garten-& Rasenkalk, Bauhaus), quartz sand (0.4 to 0.8 mm in diameter, Carl Roth GmbH) and distilled water, sealed with surgical tape (Micropore, 3M). Algae were kept under long day conditions at 22°C in a 16 h light: 8 h dark cycle with white light lateral illumination (30 to 70 µmol photons m^-2^ s^-1^, Lumilux L36W/840, OSRAM).

Algae were cut along their thalli below and above each node at the University of Marburg using razor blades and a stereomicroscope (Leica DM6000 CS). Harvested nodes were pooled, collected in 1.5 ml reaction tubes, frozen in liquid nitrogen and stored at -80°C until further analysis.

### Preparation of total RNA and Northern hybridizations

Total RNA was extracted utilizing a modified acid guanidinium thiocyanate-phenol-chloroform protocol (Chomczynski and Sacchi, 1987, 2006) using PGTX (Pinto et al., 2009), but omitting Triton X-100.

At least 100 mg FW total *Chara braunii* samples were ground into a homogenate using mortar and pestle under liquid nitrogen. Samples were suspended in 1 ml Z6 buffer (8 M guanidiniumhydrochloride, 20 mM MES, 20 mM EDTA, 50 mM 2-mercaptoethanol, pH 7), mortar rinsed with 500 µl Z6 buffer, and transferred to a screw-cap tube. 3 ml PGTX was added and samples incubated for 30 minutes at room temperature while vortexing every 5 minutes. Two chloroform extractions were performed consecutively; each adding 3 ml chloroform/isoamylalcohol (24:1), incubating samples for 10 min at room temperature under occasional vortexing, centrifugation for 3 min (3.273 g, at room temperature) and transferring the aqueous layer to a fresh tube. After chloroform extractions, RNA was precipitated using one volume 2-propanol over night at -20°C. RNA was collected by centrifugation for 30 min (13.000 g, 4°C), and washed two times using 70% ethanol and air dried for 10 minutes. RNA was resuspended in 100 µl RNase-free H_2_O and stored at -80°C until further analysis.

Nodal RNA was extracted analogous. Pooled nodal cells were suspended in 250 µl Z6 buffer in ice cooled 2 ml reaction tubes. Cell disruption was performed using a mixer mill (MM 400, Retsch; 5 cycles of 90 s with varying frequencies: 30 s 8/s, 30 s 16/s, 30 s 25/s, 2 min cooled on ice) using one glass bead (2.85 to 3.45 mm diameter, Carl Roth GmbH). Supernatant was transferred into a new reaction tube alongside 125 µl Z6 buffer rinse of glass bead. 750 µl PGTX was added and samples incubated for 30 minutes at room temperature while vortexing every 5 minutes. Addition of 750 µl chloroform/isoamylalcohol (24:1) was followed by incubation for 10 min at room temperature under occasional vortexing. Afterwards, sample tubes were centrifuged for 3 minutes (room temperature, 3273 g). Following another chloroform extraction, the upper aqueous phase was precipitated with one volume 2-propanol over night at -20°C. RNA was collected by centrifugation for 30 min (13.000 g, 4°C) and air dried for 10 minutes. RNA was resuspended in 20 µl RNase-free H_2_O and stored at -80°C until further analysis.

RNA concentration and purity were measured using a NanoDrop ND-1000 spectrophotometer according to manufacturer instructions (PEQLAB Biotechnologie GmbH). Residual DNA was removed using the Turbo DNase-free™ Kit (Thermo Fisher Scientific, Germany) following manufacturer instructions. RNA was recovered using RNA Clean & Concentrator kits (Zymo Research, USA). RNA integrity was controlled on a 5200 Fragment Analyzer System, according to manufacturer instructions (Agilent, USA).

8 µg total RNA was separated by 1.4% denaturing agarose gel electrophoresis and transferred by capillary transfer on Hybond N+ nylon membranes (Cytiva Europe GmbH) over night. Northern hybridization was performed with radioactively labelled probe for chlorophyll a/b binding protein (*cab*) mRNA (primers for template; see **Table 1** for sequences), generated using [_α_-32P]-UTP and the Maxiscript T7 *in vitro* transcription kit (Thermo Fisher Scientific, Germany). Blotted RNA was crosslinked to the membrane via 240 mJ using a UV-Stratalinker (Stratagene). Hybridizations were performed in Northern buffer (50% deionized formamide, 7% SDS, 250 mM NaCl and 120 mM Na_2_HPO_4_/NaH_2_PO_4_ pH 7.2) overnight at 62°C. The membranes were washed at 57°C in buffer 1 (2×SSC (3 M NaCl, 0.3 M sodium citrate, pH 7.0), 1% SDS), buffer 2 (1×SSC, 0.5% SDS) and buffer 3 (0.1×SSC, 0.1% SDS) for 10 min, 5 min and 2 min. Hybridization signals were detected via Typhoon FLA 9500 imaging system (GE Healthcare, USA) using phosphorimaging and quantified using Quantity One software (Bio-Read Laboratories Inc., USA).

**Table 1.**
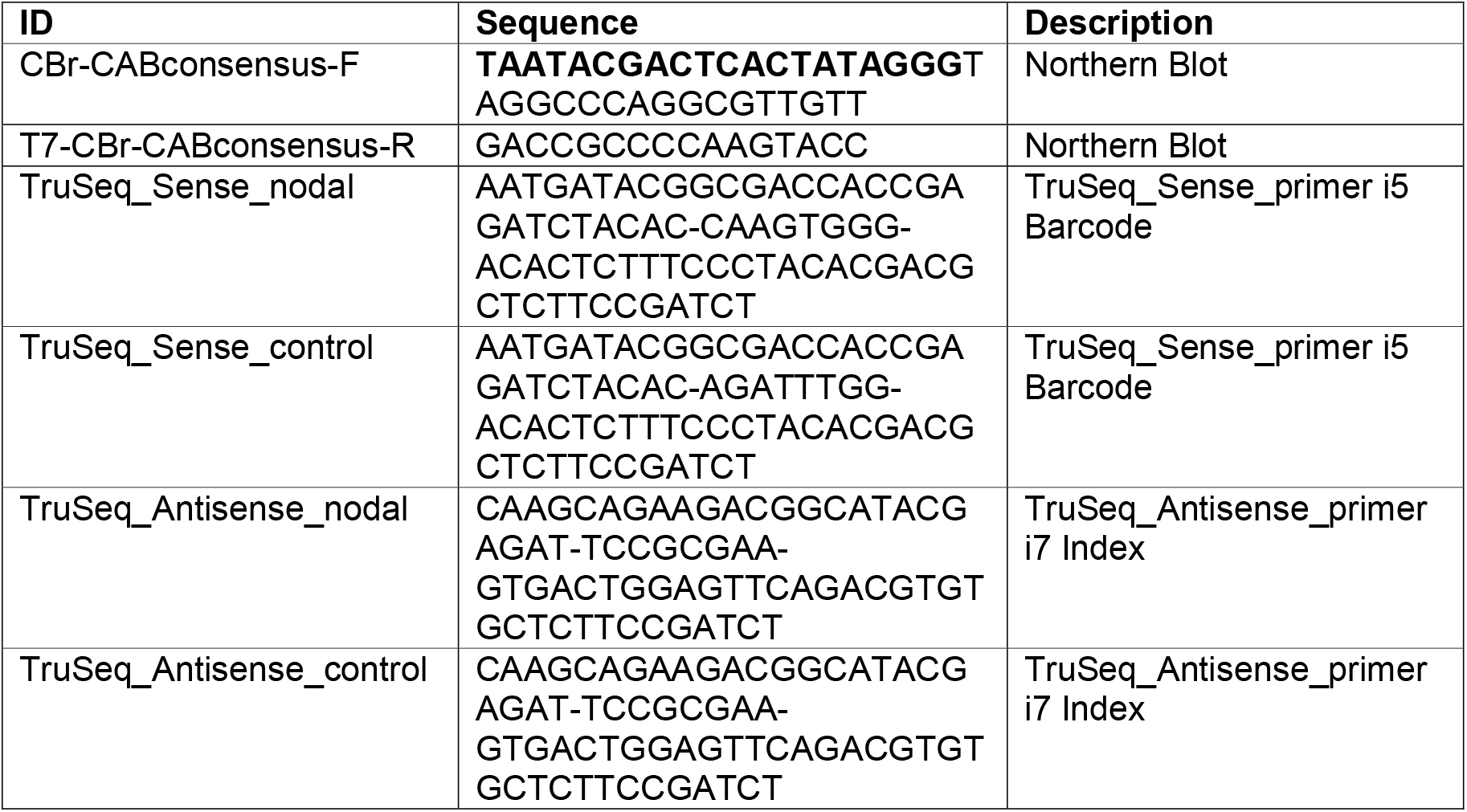
Oligonucleotides used in this study. An added T7 promoter sequence is highlighted in boldface letters.

Templates were amplified from *Chara* genomic DNA using OneTaq® Quick-Load® 2X Master Mix with Standard Buffer. To facilitate the use of PCR products as templates for *in vitro* transcription, the T7 RNA polymerase promoter (TAATACGACTCACTATAGGG) was included in the 5’ sections of the corresponding reverse primers (**Table 1)**.

### Preparation of total DNA from *Chara braunii*

Genomic DNA was extracted from at least 100 mg FW total *Chara braunii* samples ground into a homogenate using mortar and pestle under liquid nitrogen. Samples were suspended in 1 ml SET buffer (1 mM EDTA, 25% sucrose, 50 mM Tris) and lysed over night at 50°C in a screw-cap tube after adding 1/4 volume 0.5 M EDTA, 1/10 volume 20% SDS and 100 µg/ml proteinase K (Scholz et al., 2019).

After the addition of 2 volumes of ROTI® Aqua-Phenol (DNA; Carl Roth GmbH, Germany) and 2 volumes of chloroform/isoamylalcohol (24:1), samples were incubated for 5 minutes at room temperature while vortexing intermediately. Samples were centrifuged for 5 min (3273 g, room temperature), the aqueous layer transferred to a fresh tube and the chloroform extraction step repeated. Afterwards, DNA was precipitated using one volume 2-propanol over night at -20°C, collected by centrifugation for 30 min (13.000 g, 4°C), and washed two times using 70% ethanol and air dried for 10 minutes. DNA was resuspended in 100 µl RNase-free H_2_O and stored at -20°C until further analysis.

### cDNA sequencing

cDNA libraries were constructed and sequenced as a service provided by vertis Biotechnologie AG (Germany). Poly(A)+ RNAs were fragmented using ultrasound (one pulse of 30 s at 4°C), after which an oligonucleotide adapter was ligated to the 3’ termini. First-strand cDNA synthesis was performed using M-MLV reverse transcriptase and the 3’ adapter as primer. First strand cDNA was purified and the 5’ Illumina TruSeq sequencing adapter ligated to the 3’ end of the antisense cDNA. Resulting cDNA was PCR-amplified using a high fidelity DNA polymerase. The cDNA was purified using an Agencourt AMPure XP kit (Beckman Coulter Genomics, USA). For Illumina NextSeq sequencing, samples were pooled in approximately equimolar amounts, purified using the Agencourt AMPure XP kit (Beckman Coulter Genomics, USA) and analyzed by capillary electrophoresis. Primers used for PCR amplification were designed for TruSeq sequencing according to manufacturer instructions. The cDNA pool was single-read sequenced on an Illumina NextSeq 500 system using 75 bp read length.

### Bioinformatic analyses

Data analysis was performed on the European instance galaxy server (The Galaxy Community, 2022) following the available guidelines for reference based RNA-seq analysis (Batut et al., 2018, 2022). The quality of raw reads was assessed using FastQC (Andrews, 2010) and MultiQC (Ewels et al., 2016). Trimming of adapter and barcode contamination was performed using cutadapt (Martin, 2011). Mapping and counting of reads against the published *Chara* genome (Nishiyama et al., 2018) was performed via RNA STAR (Dobin et al., 2013). Differential gene expression analysis was performed via manual log2FC calculation following the given guidelines.Differentially expressed genes were determined with a cutoff as log_2_FC ≥ │1│. Gene ontology (GO) analysis of RNA-seq data was performed using goseq (Galaxy version 1.44.0) (Young et al., 2010).

### Phylogenetic Analysis

A set of amino acid sequences (**Supplementary Table S1**) was selected based on position within the green lineage and sequence similarity to *Chara braunii* putative amylase homologous genes of interest, g31563 and g41182. The sequences were aligned with M-coffee (Notredame et al., 2000; Di Tommaso et al., 2011) and further analysed using the BEAST 2 software package version 2.7.3 (Bouckaert et al., 2019). Calculations were performed using standard BEAUTi settings utilizing the tree prior yule model, substitution model Blosum62 and MCMC chain length of 1e6, with logged parameters every 1e4 steps. Tracer v1.7.2 software (Rambaut et al., 2018) was used to validate the effective sample sizes (ESS>200). TreeAnnotator was used to build maximum clade credibility trees using burnIn of 50% and a posterior probability limit of 0.5 for median node heights. FigTree v1.4.4 software (Rambaut, 2018) was used to visualise the generated dendrograms.

## RESULTS AND DISCUSSION

### The most abundantly transcribed genes in enriched *Chara* nodal cells

The nodal cells of *Chara braunii* are embedded in the zone between two adjacent large internodal cells from where the branchlet and stipulopode cells are emanating (**Figure 1**). The algae were cut under the stereomicroscope along their thalli below and above each node, branchlets were removed near the node. Separated nodes were kept in liquid nitrogen until RNA extraction. RNA was extracted from 28 pooled nodes, yielding 0.68 µg of total RNA after DNase treatment. In parallel, RNA was extracted from fresh total algal material. Judged by the visible sharp rRNA bands and the test hybridization for *cab* mRNA, the prepared RNA samples were of high quality (**Figure 2**). After polyA-mRNA enrichment, cDNA libraries were generated for the two different *Chara braunii* RNA samples and sequenced yielding a total of 54.042.636 raw single read counts for enriched nodal cells and 53.423.073 for total cells. From these, 10.991.291 (20.6%) for total, and 13.763.747 (25.5%) reads for the enriched sample remained unmapped. After trimming and mapping steps, 4.994.360 nodal (9.4%) and 2.628.740 total (4.9%) reads were uniquely mapped to the *Chara* genome (**Figure 3**). Mapped reads matched 11.319 putative genes for the nodal sample, and 15.627 genes for total sample.

**FIGURE 1.**
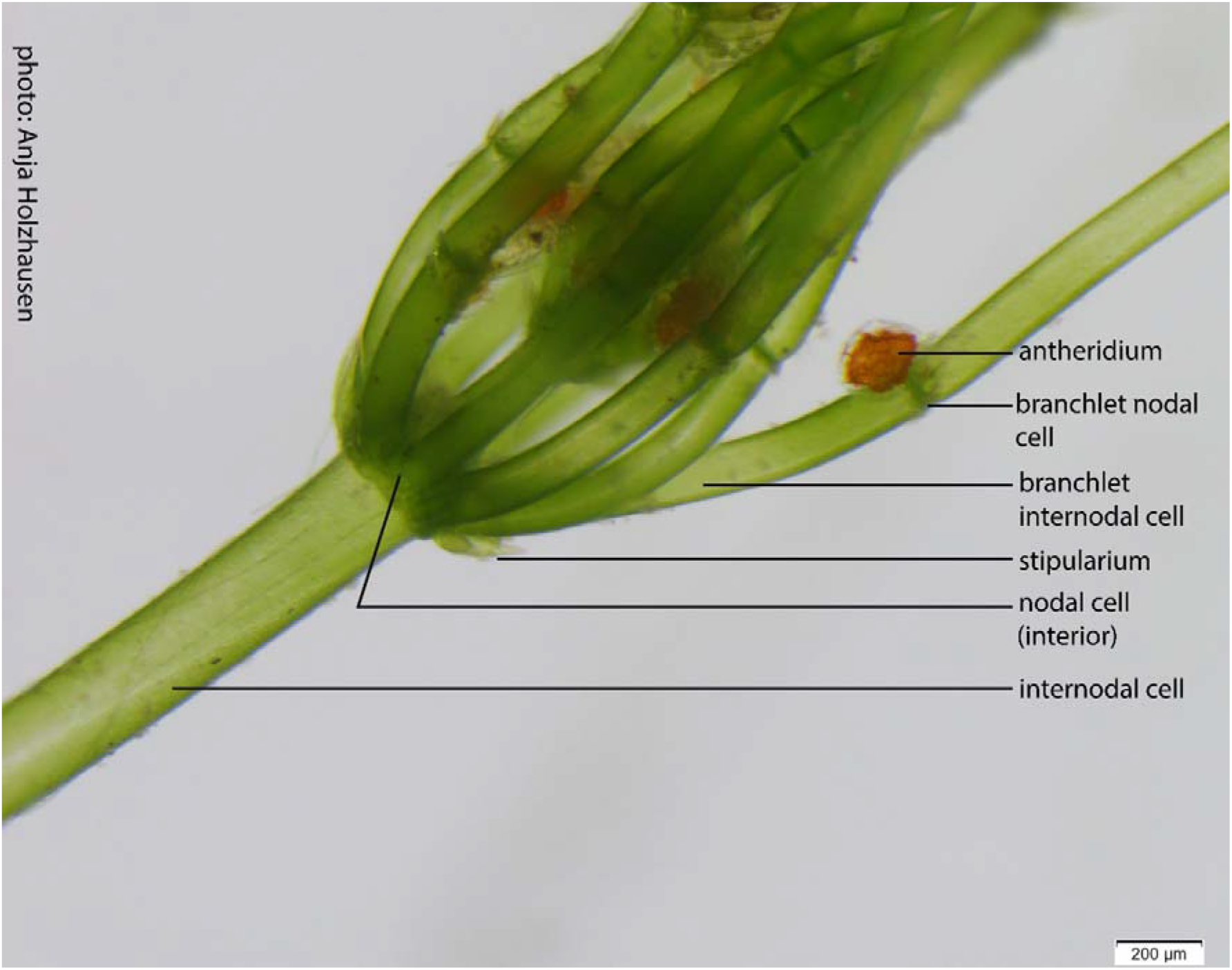
Specimen of *Chara baunii*. Detailed view of *Chara braunii* S276. Shown is the area with the interior node cells confined by two internodal cells. Nodal cells pinched off peripheric cells from which branchlets developed with a structure analogous to the main stem consisting of branchlet nodal and internodal cells. Branchlet nodal cells are the origin for the development of antheridia (shown) and the typical charophytic oogonia after proliferation.

**FIGURE 2.**
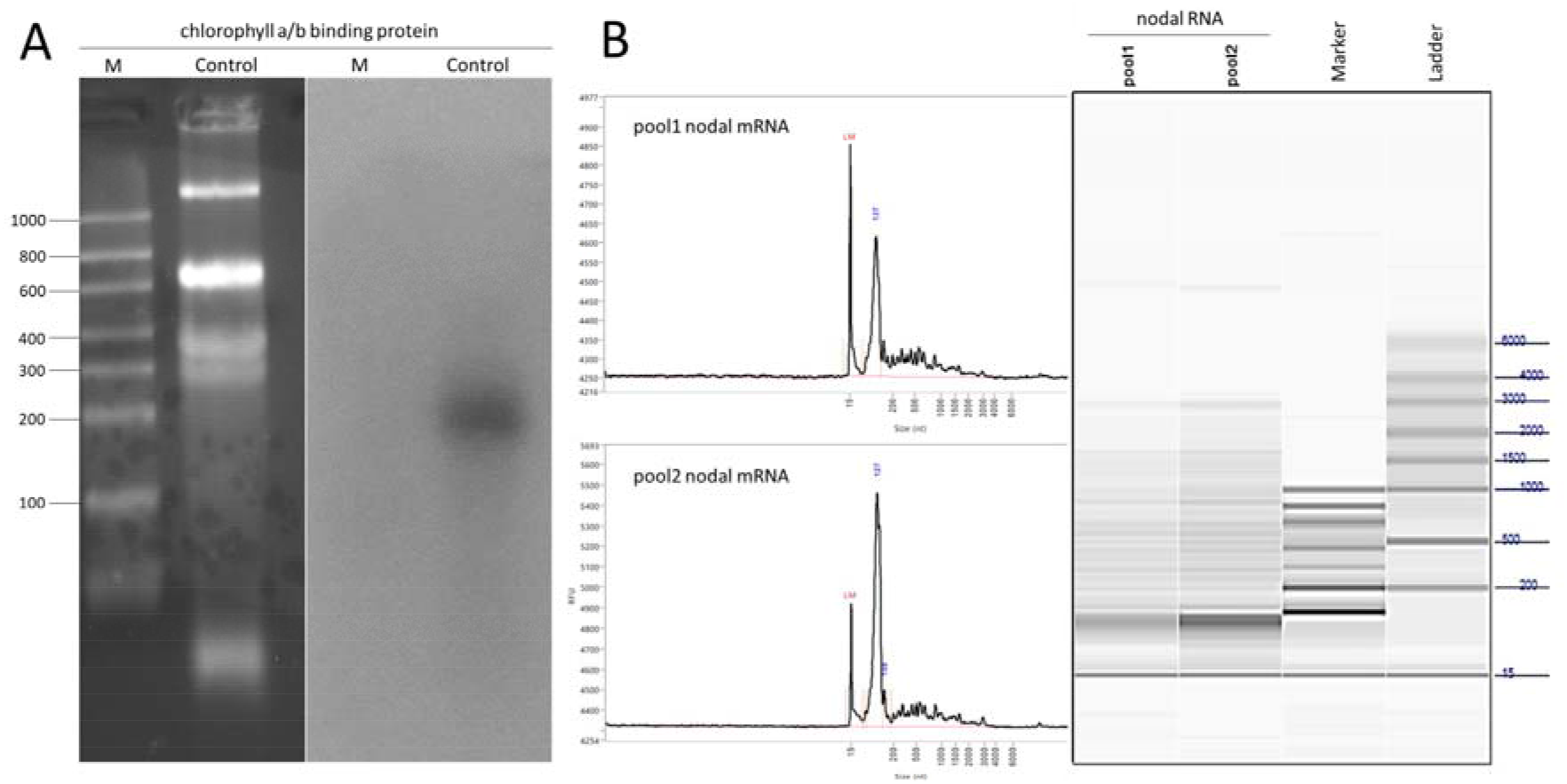
*Chara braunii* RNA samples. **(A)** Northern blot analysis of *Chara braunii cab*. 8 µg total RNA was run on a 1.4% denaturing agarose gel (left). Clear appearance of bands without smearing indicates high quality of RNA. The membrane was probed with ^32^P-labeled *cab* single-stranded RNA probe (right). M = Ribo Ruler Low Range, Control = total *Chara* RNA.**(B)** Fragment analyzer run and electropherograms of pooled, enriched *Chara braunii* nodal RNA after DNase treatment. Distinct appearance of bands without smearing indicates low degradation of RNA. Samples include 15 nt internal marker. Marker = Ribo Ruler Low Range, Ladder = HS RNA ladder.

**FIGURE 3.**
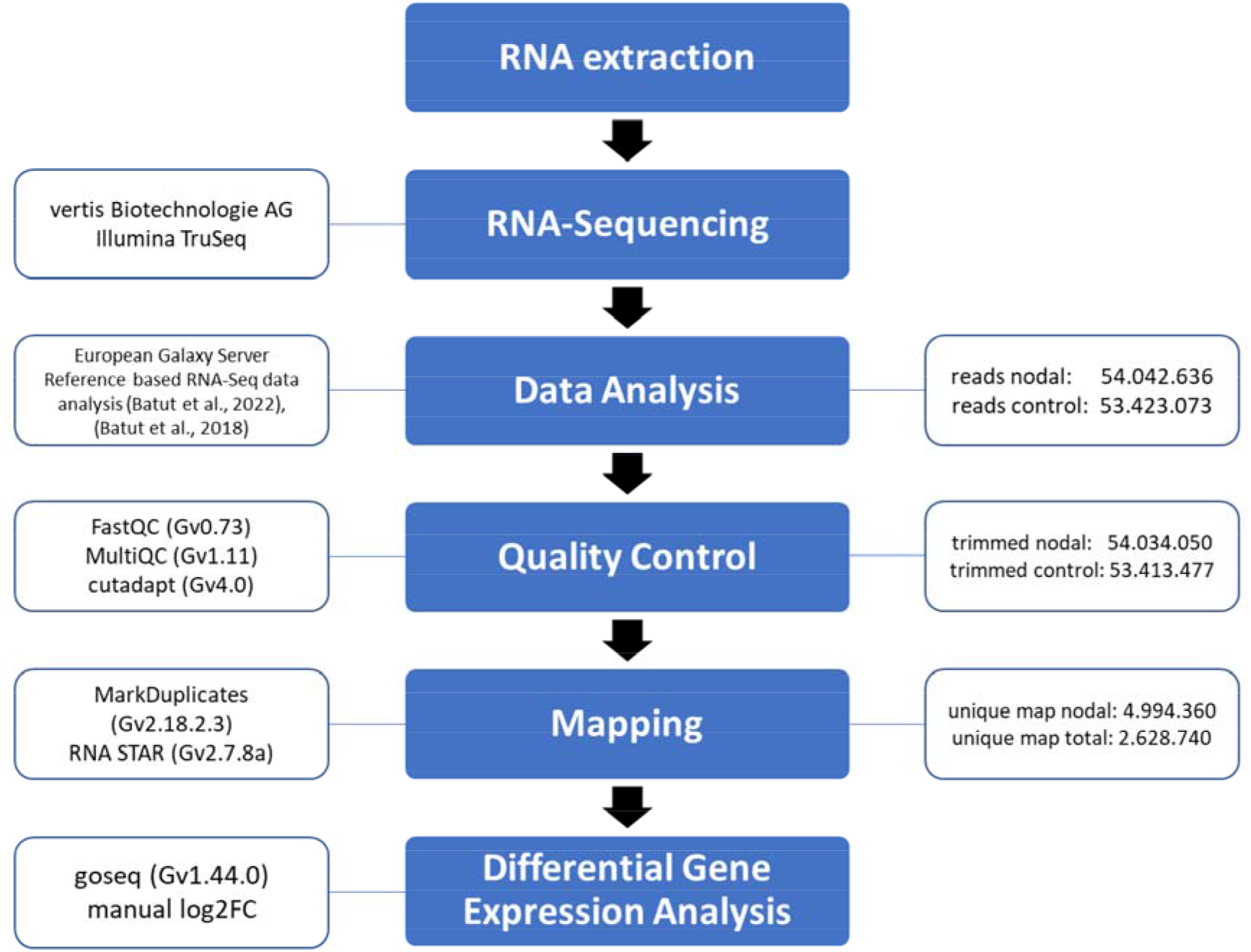
RNA-seq data analysis workflow. Overview of steps performed for RNA-seq analysis of *Chara braunii* transcriptome. Blue boxes (middle) illustrate steps within the workflow from RNA extraction to gene expression analysis, transparent boxes (left) illustrate the methods and bioinformatic tools used, transparent boxes (right) illustrate the number of reads at various points of the bioinformatic analysis of transcriptome data. Data analysis was performed on the European galaxy server (The Galaxy Community, 2022), based on the published guidelines for reference based RNA-seq data analysis (Batut et al., 2018, 2022).

The 15 most highly expressed genes for each tissue sample are given in **Tables 2** and **3**. The most abundantly expressed genes in the control sample were mostly associated with photosynthesis and protein synthesis, such as chlorophyll a-b binding proteins (5/15 instances), the photosystem (PS) II 5 kDa protein PsbT, or the ribosomal proteins RpL6-like, RpL35a-3-like, RpL10, RpS19-3 and RpS8 (**Table 2**). Some of the ribosomal proteins encoded by the detected mRNAs are universally found throughout the tree of life (Lan et al., 2022), and some facilitate also extraribosomal functions (reviewed in Xiong et al., 2021). RpL6 has been shown to promote growth, productivity and tolerance towards salt stress in rice (Moin et al., 2021). RpL35a in humans has been shown to inhibit cell death (Lopez et al., 2002). Finally, RpL10 is not only highly conserved due to its critical function in joining the 40S and 60S ribosomal subunits into a functional 80S ribosome, but also involved in cell growth and resistance to non-host disease and reactive oxygen species in plants (Ferreyra et al., 2010; Ramu et al., 2020).

**Table 2.** Most abundant mRNAs in control. The 15 genes for which the highest read counts in the control sample were detected in descending order. The gene ID in the first column is followed by the annotation and the normalized read counts in the control and in the nodal samples.

| Gene ID | Description | control counts | nodal counts |
| --- | --- | --- | --- |
| g49772 | n/a | 820153 | 1875 |
| g50349 | chlorophyll a-b binding protein | 76820 | 7262 |
| g49773 | n/a | 74655 | 178 |
| g48962 | n/a | 56585 | 128 |
| g20157 | chlorophyll a-b binding protein 151, chloroplast | 53471 | 1408 |
| g18875 | PSII 5 kDa PsbT protein, chloroplast | 50667 | 5196 |
| g20098 | 60S ribosomal L6-like protein | 39190 | 38394 |
| g20154 | major chlorophyll binding protein | 37336 | 2194 |
| g20151 | chlorophyll a-b binding protein 151, chloroplast | 36381 | 740 |
| g50044 | 60S ribosomal protein L35a-3-like | 33211 | 20144 |
| g29937 | 60S ribosomal protein L10 | 29890 | 21826 |
| g21828 | 40S ribosomal protein S19-3 | 29726 | 19115 |
| g32357 | 40S ribosomal protein S8 | 29627 | 18184 |
| g19118 | chlorophyll a-b binding protein 151, chloroplast | 26204 | 1230 |
| g46838 | PAX3- and PAX7-binding protein 1 | 25206 | 30172 |

The high expression of genes encoding photosynthesis- and ribosome-associated proteins is consistent with the photosynthetic lifestyle and fast growth of the *Chara braunii* algae. Another three of these mRNAs encode unknown proteins, while with g46838 also an mRNA encoding a GC-rich sequence DNA-binding factor-like protein with similarity to the PAX3- and PAX7-binding proteins was found. The homologs of the latter in animals are involved in epigenetic regulation (Diao et al., 2012), but there is no information for putative plant homologs.

The list of the 15 most strongly expressed genes in the sample enriched for nodal cells differed from the control sample by lacking all genes related to photosynthesis. However, even 9/15 mRNAs were encoding ribosomal proteins indicating that the drive for protein synthesis was high in this tissue (**Table 3**). Another four mRNAs encode unknown proteins, while with g26356 a thioredoxin-encoding mRNA and with g46838 again the mRNA encoding a putative GC-rich sequence DNA-binding factor-like protein with similarity to the PAX3- and PAX7-binding proteins was found.

**Table 3.** Most abundant mRNAs in central and nodal cells. The 15 genes for which the highest read counts in the nodal sample were detected, in descending order. The gene ID in the first column is followed by the annotation and the normalized read counts in the nodal and in the control samples.

| Gene ID | Description | nodal counts | control counts |
| --- | --- | --- | --- |
| g20098 | 60S ribosomal L6-like protein | 38394 | 39190 |
| g46838 | PAX3- and PAX7-binding protein 1 | 30172 | 25206 |
| g29937 | 60S ribosomal protein L10 | 21826 | 29890 |
| g10845 | n/a | 20269 | 19117 |
| g50044 | 60S ribosomal protein L35a-3-like | 20144 | 33211 |
| g21828 | 40S ribosomal protein S19-3 | 19115 | 29726 |
| g17012 | ribosomal protein L3, conserved site | 18638 | 20362 |
| g32357 | 40S ribosomal protein S8 | 18184 | 29627 |
| g41408 | 60S ribosomal protein L5 | 13381 | 23292 |
| g26356 | thioredoxin H-type | 13127 | 6048 |
| g12280 | n/a | 13114 | 9764 |
| g38728 | n/a | 13063 | 22161 |
| g23412 | 60S ribosomal protein L17-2-like | 12797 | 25171 |
| g8281 | 40S ribosomal protein S21-2 | 11504 | 10917 |
| g36718 | n/a | 10474 | 5103 |

The full list of genes for which mRNAs were detected is given in **Supplementary Table S2**. Interestingly, 117 different mRNAs for proteins with domains of unknown function (DUF) were found as well. Among them, we found 21 instances of genes encoding DUF4360*-*containing cysteine-rich proteins. While DUF4360*-*containing proteins exist in bacteria and eukaryotes (Paysan-Lafosse et al., 2023), it is striking that these *Chara braunii* DUF4360*-*containing proteins are more similar to homologs from fungi (mainly Ascomycota) than to any other. Most of the mRNAs for DUF4360*-* containing proteins were more abundant in the *Chara* total control sample (e.g., genes g9217, g68682, g2717, g2719, g58375, g11068, g36724, g68700, g9208, g68689, g11072 and g76151; for details see **Supplementary Table S2**).

### Functional assignments to differentially expressed genes

To obtain an overview about genes differentially expressed between the two samples, we set cut-offs of log_2_FC ≥ │1│ and RPK ≥10 leaving a set of 3145 differentially regulated genes (2762 up, 383 down in the sample enriched for nodal cells compared to control). The full list of genes can be found in **Supplementary Table S2**.

Gene ontology (GO) assignments were used to classify the putative functions of the differently expressed *Chara braunii* genes (**Figure 4**). We sorted the GO assignments according to biological process, cellular components and molecular function, and into the up-or down-regulated categories. Among the upregulated biological processes in the nodal sample were the GO terms “protein folding” (5 genes), “lipid metabolic process” (4 genes), “transmembrane transport” (5 genes) and “regulation of transcription, DNA-templated” (5 genes) most prevalent. Prominent cellular component GO terms were “integral component of membrane” (17 genes), “membrane” (12 genes) and “nucleus” (8 genes). Among GO terms for molecular function, “protein binding” (223 genes), “nucleic acid binding” (59 genes), “binding” (39 genes) and “DNA binding” (32 genes) were prevalent.

**FIGURE 4.**
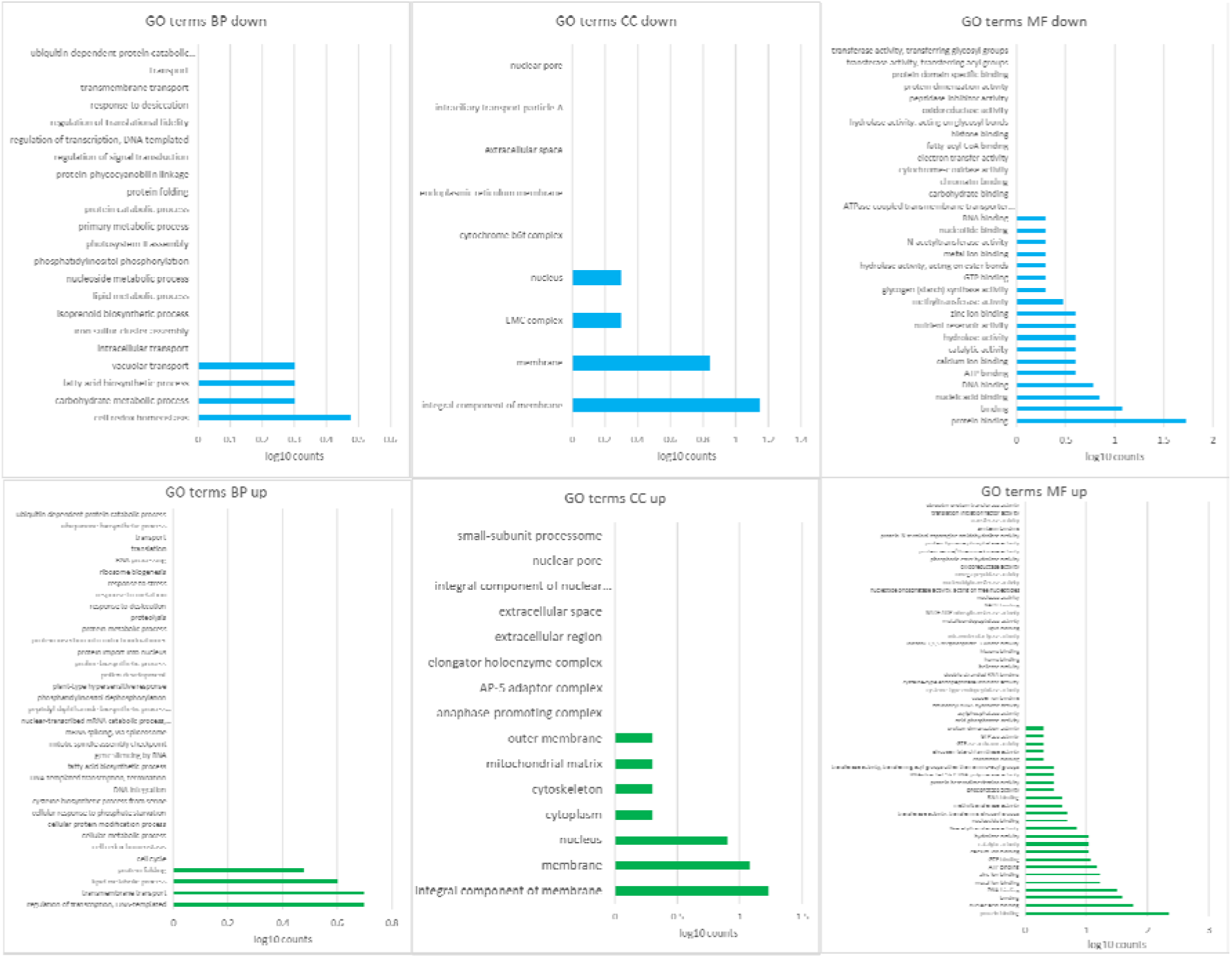
GO term analysis of enriched *Chara* central and nodal cells. Chart representations of gene ontology terms enriched in nodal and central cells compared to total *Chara braunii* within the log_2_FC ≥ │1│ dataset. GO terms for biological processes (BP), cellular components (CC) and molecular function (MF) are shown as log10 transformed counts, separated by up-and downregulation. Green bar terms indicate upregulation, blue bar terms indicate depletion.

In contrast, the biological process GO terms “cell redox homeostasis” (3 genes), “carbohydrate metabolic process” (2 genes), “fatty acid biosynthetic process” (2 genes) and “vacuolar transport” (2 genes) were downregulated. Among downregulated cellular component GO terms, “integral component of membrane” (14 genes), “membrane” (7 genes), “EMC complex” (2 genes) and “nucleus” (2 genes) were most prevalent. Prominent GO terms for downregulated molecular functions were “protein binding” (53 genes), “binding” (12 genes), “nucleic acid binding” (7 genes) and “DNA binding” (6 genes).

Because there were many genes with unclear annotation or similarity to retrotransposons, we will in the following focus on genes with an available functional annotation (**Table 4**).

**Table 4.** Selected up-and down-regulated genes in nodal cells compared to the control mentioned in the text. The gene ID in the first column is followed by the annotation and the calculated differential expression (log_2_FC). A positive value for the log_2_FC indicates higher read count in the nodal sample (shaded in light brown), while a negative value indicates a higher read count in the control (shaded in light green).

| Gene ID | Description | $\log_2FC$ |
| --- | --- | --- |
| g31563 | putative alpha-amylase, 1,4-alpha-glucan-branching protein | 5.19 |
| g29739 | caffeoylshikimate esterase | 4.93 |
| g48048 | mediator of RNA polymerase II transcription subunit 19a-like | 4.61 |
| g19216 | Spastin | 4.59 |
| g41182 | beta-amylase 3, chloroplast | 4.38 |
| g23946 | protein TRIGALACTOSYLDIACYLGLYCEROL 3 (TGD3), chloroplast | 4.07 |
| g19355 | serine/threonine-protein kinase EDR1 isoform X1 | 2.60 |
| g55470 | serine/threonine-protein kinase EDR1-like isoform X1 | 2.59 |
| g17647 | ETHYLENE INSENSITIVE 3-like 1 protein | 2.38 |
| g197 | probable ADP-ribosylation factor GTPase-activating protein AGD11 | 2.34 |
| g84830 | START-like domain homeobox protein Wuschel-like | 2.33 |
| g30098 | ETHYLENE INSENSITIVE2 | 2.11 |
| g32224 | PAP11, MurE-like_UDP-N-acetylmuramoyl-L-alanyl-D-glutamate--2, 6-diaminopimelate ligase | 1.17 |
| g19139 | chlorophyll a-b binding protein 50, chloroplast | -1.44 |
| g51450 | PAP10, thioredoxin-like protein CITRX, chloroplast | -1.45 |
| g40841 | chlorophyll a-b binding protein CP26, chloroplast | -1.91 |
| g20154 | major chlorophyll binding protein | -2.10 |
| g72716 | chlorophyll a-b binding protein P4, chloroplast | -2.27 |
| g19118 | chlorophyll a-b binding protein 151, chloroplast | -2.43 |
| g17032 | PAP9, Superoxide Dismutase, FeSOD | -2.57 |
| g19140 | chlorophyll a-b binding protein 50, chloroplast | -2.58 |
| g31512 | photosystem I reaction center subunit VI, chloroplast | -2.89 |
| g19133 | major chlorophyll binding protein | -3.20 |
| g20157 | chlorophyll a-b binding protein 151, chloroplast | -3.26 |
| g50764 | photosystem II repair protein PSB27-H1, chloroplast | -3.30 |
| g61495 | peroxiredoxin-2B | -3.40 |
| g48045 | PAP5 | -3.63 |
| g20151 | chlorophyll a-b binding protein 151, chloroplast | -3.64 |
| g602 | protein LHCP TRANSLOCATION DEFECT | -3.93 |
| g19141 | chlorophyll a-b binding protein 3C, chloroplast | -5.16 |
| g20155 | major chlorophyll binding protein | -5.37 |
| g19126 | chlorophyll a-b binding protein 50, chloroplast | -6.79 |

Among the genes with higher expression in nodal cells are two, g31563 (log_2_FC 5.19) and g41182 (log_2_FC 4.38), that encode proteins with similarity to 1,4-alpha-glucan-branching enzymes/alpha/maltogenic amylase and beta amylase. While it is known that charophytes use starch as a storage product (García, 1994), the upregulation of both genes in the nodal cell sample indicates active carbohydrate storage molecule metabolism in the respective cells of *Chara braunii*.

With their publication of the *Chara* draft genome, Nishiyama et al. (2018) predicted the presence of certain factors involved in phytohormone biosynthesis and signalling pathways. Within our dataset, we detected several upregulated genes belonging to these pathways. Related to ethylene signaling, we detected three upregulated genes in the nodal samples. The genes g30098 (log_2_FC+2.1) and g17647 (log_2_FC+2.4) encode putative homologs of the ETHYLENE INSENSITIVE 2 and 3 proteins (EIN2 and EIN3) (Nishiyama et al., 2018), proteins that are central in the regulation of ethylene-mediated responses (Chao et al., 1997; Solano et al., 1998). Furthermore, we detected putative serine/threonine-protein kinases EDR1, g19355 (log_2_FC+2.6) and g55470 (log_2_FC+2.6), homologs of two proteins which in *A. thaliana* are involved in crosstalk between ethylene-, ABA- and salicylic acid signaling (Tang et al., 2005; Wawrzynska et al., 2008). Additionally, we detected with a log_2_FC of +4.1 high differential expression of g23946, encoding a homolog of the trigalactosyldiacylglycerol 3 (TGD3) protein, involved in chloroplast lipid import (Lu et al., 2007), jasmonic acid signalling and plant defence responses via phosphatidic acid signalling (Tagami et al., 2021).

Regarding further upregulated, signalling-related genes, we detected with g197 (log_2_FC+2.3) an mRNA encoding a putative ADP-ribosylation factor GTPase-activating protein (ARF). Another upregulated gene, g48048 (log_2_FC+4.6), encoding a putative mediator of RNA polymerase II transcription subunit 19a-like, points towards an increase in transcriptional activity for the actively dividing nodal cells. Complementary, g19216 (log_2_FC+4.6), a putative spastin-like protein, has been stated to coordinate microtubule and endoplasmatic reticulum (ER) dynamics. While spastins modulate the lipid droplet network and their dispersion throughout the cell in animals, this has been speculated to be an ancestral process in eukaryotes, as antecedent ER elongation along microtubules is a conserved mechanism also described in plants (Hamada et al., 2014; Arribat et al., 2020).

Finally, we detected g29739 (log_2_FC+4.9), a putative caffeoylshikimate esterase (CSE) involved in the biosynthesis of lignin, which was already confirmed to be present in *Chara* through biochemical analysis (Rekha and Sujathamma, 2020).

Within our dataset, we detected several genes encoding proteins involved in photosynthetic processes which were expressed at lower levels in nodal cells. Among them, the genes g19126, g20155, g19141, g20151, g20157 and g19133 (log_2_FC -6.8, -5.4, -5.2, -3.6, -3.3, -3.2) encode chlorophyll a-b binding proteins, apoproteins of the light-harvesting complex of photosystem II (Liu et al., 2013). Additionally, g31512 (log_2_FC -2.9), encoding the PSI reaction center subunit VI, g50764 (log_2_FC -3.3) encoding the putative PSII repair protein PSB27 and g602 (log_2_FC -3.9) encoding a putative LHCP translocation defect protein, point at less active photosynthetic processes in central and nodal cells. Interestingly, g61495 (log_2_FC -3.4), a putative peroxiredoxin involved in cellular protection from reactive oxygen species (ROS), was downregulated as well. Besides their crucial scavenging capabilities, however, peroxiredoxins function also as modulators of stress response pathways in their interaction with transcription factors (Bréhélin et al., 2003; Hopkins and Neumann, 2019).

Another set of proteins with central functions in the chloroplast are the PEP-associated proteins (PAPs) which control the activity of the plastid-encoded RNA polymerase (PEP) (Pfalz and Pfannschmidt, 2013). In land plants such as *Arabidopis thaliana*, 12 PAPs were described (Pfalz and Pfannschmidt, 2013). These PAPs are associated with the development of chloroplasts in photosynthetically active cells and therefore constitute highly selective cell type-specific markers. Ten PAP orthologs were previously detected in *C. braunii* in contrast to only 5 or 8 in *C. reinhardtii* and *Klebsormidium nitens*, respectively (Nishiyama et al., 2018). Here, we detected a lower expression for several of these PAPs in the nodal sample compared to the control, with log_2_FC of -3.63, -2.57 and -1.45 for PAP5, PAP9 and PAP10 (genes g48045, g17032 and g51450), which hence is consistent with their role in photosynthetically active cells. However, we detected for one of these proteins, the MurE-like PAP11 (gene g32224) a higher expression in the nodal material (log_2_FC +1.17). This might be related to the suggestion that algae, similar to findings in mosses (Garcia et al., 2008) encode a different, non-PAP version of this MurE-like protein (Nishiyama et al., 2018).

### Divergent phylogenies for paralogous genes highlight evolutionary innovations

The putative 1,4-alpha-glucan-branching enyzme/alpha amylase g31563 and the beta amylase g41182 highly activated in nodal cells also showed an interesting phylogenetic divergence (**Figures 5 and 6**). g31563 encodes a protein with high sequence conservation to homologs from other algae (51% identical and 68% similar amino acids in a 572 residues-long overlap with the homolog in the Klebsormidiophyceae alga *K. nitens* (accession no. GAQ89829) and 47% identical and 65% similar amino acids in a 585 residues-long overlap with the homolog in the Zygnemophyceae algae *Closterium* sp. NIES-68 (accession no. GJP44716). The next-closely related proteins are mainly from Apicomplexa such as *Toxoplasma gondii*, and other Alveolata, the Chlamydomonadales and other green algae.

**FIGURE 5.**
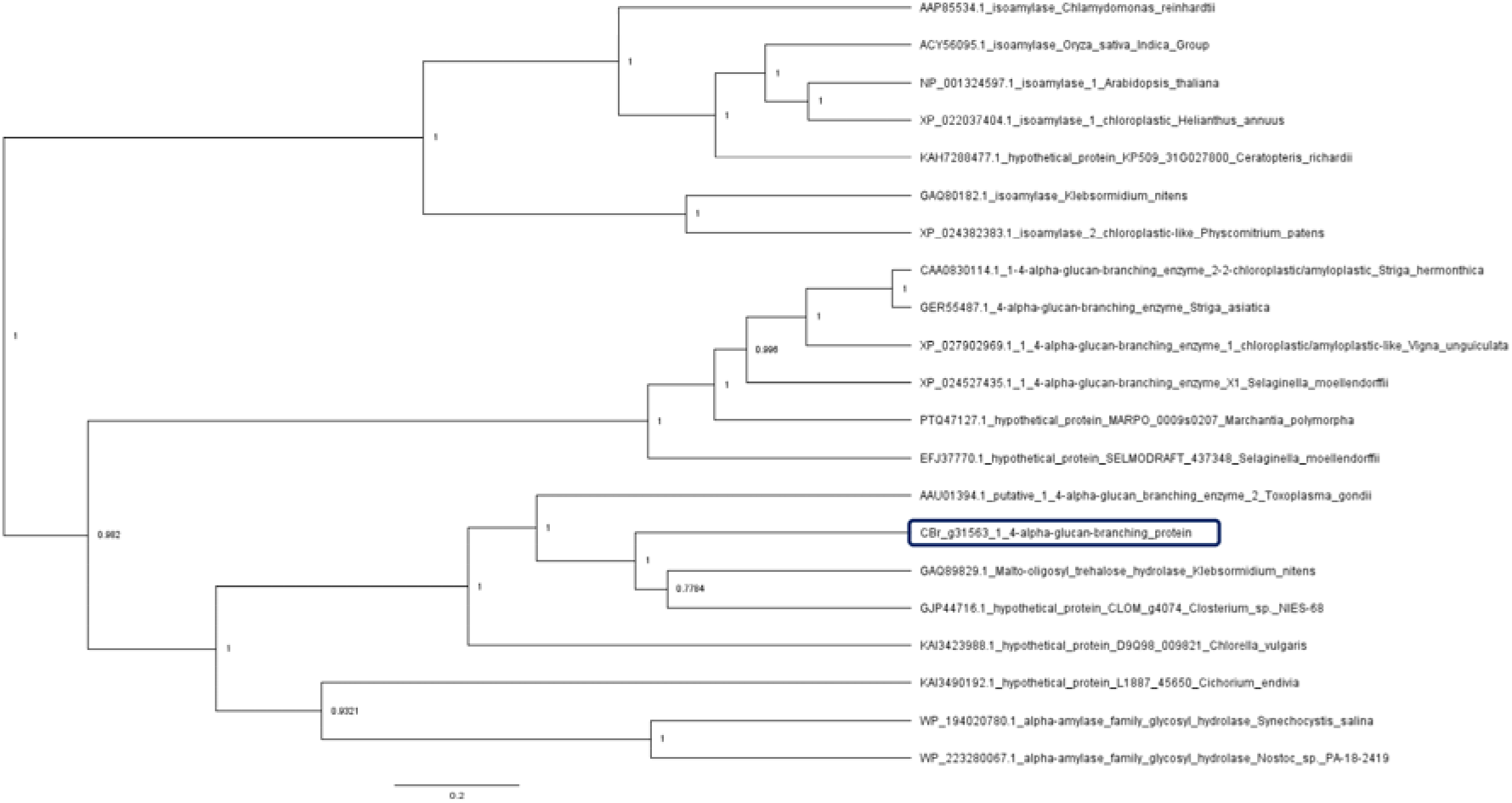
Phylogenetic analysis of a *Chara braunii* protein with similarity to 1,4-alpha-glucan-branching enzymes and alpha amylases. Bayesian inference of *Chara braunii* g31563 protein. Identified homologous proteins were aligned via M-Coffee and analyzed using BEAST 2. The depicted maximum clade credibility tree, generated via tree prior yule model and 1e4 logged MCMC chain length of 1e6, was built using TreeAnnotator with 50% burnin and a posterior probability limit of 0.5 for median node heights. The generated dendrogram was visualised using FigTree with posterior probabilities labelled on their respective nodes. The position of the putative *Chara braunii* g31563 alpha amylase homolog is boxed. Two cyanobacterial sequences (*Nostoc* sp. and *Synechocystis salina*) were added as outgroups. The compared protein sequences can be found in **Supplementary Table S1**.

**FIGURE 6.**
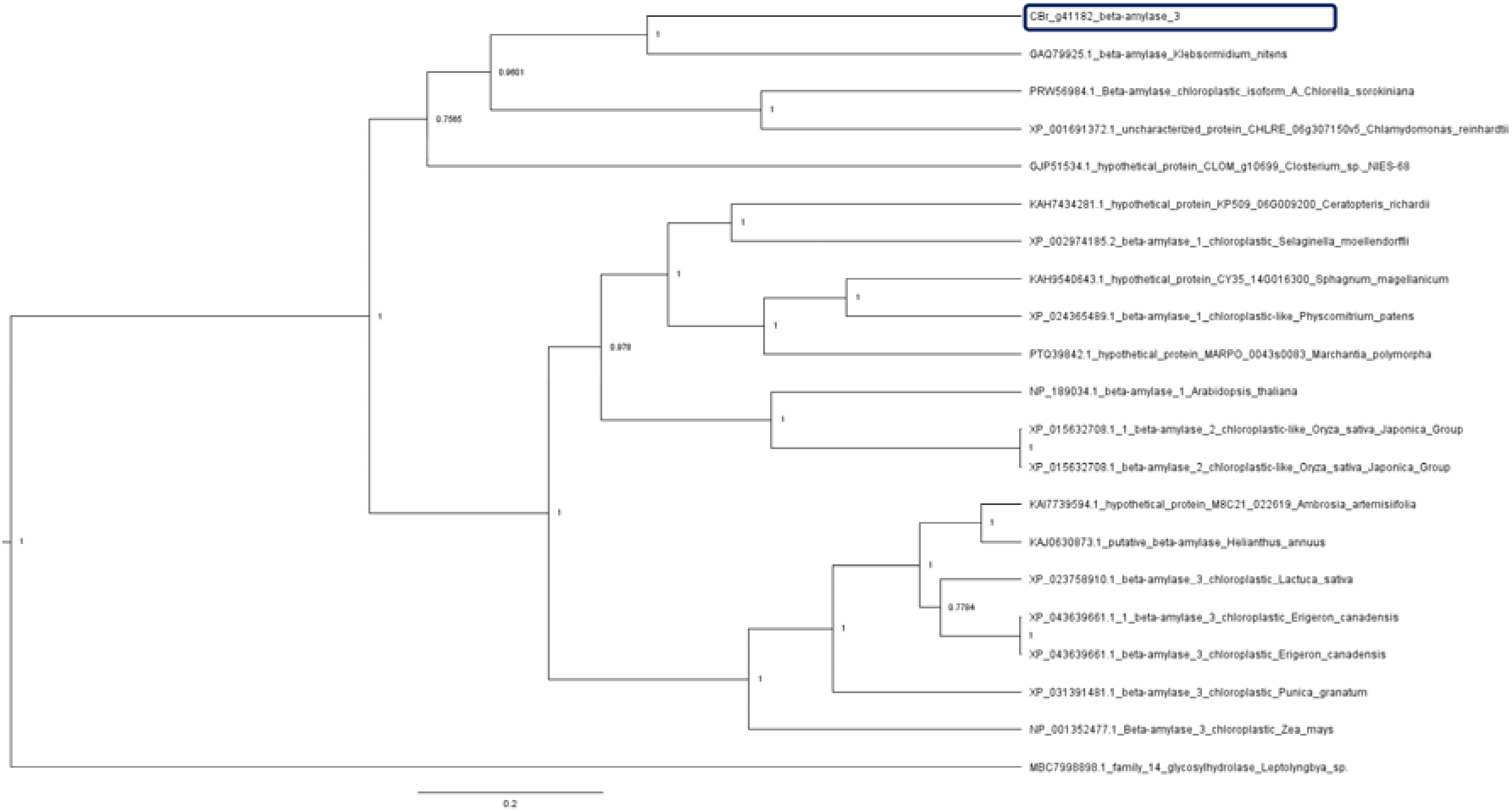
Phylogenetic analysis of a putative *Chara braunii* beta amylase homolog. Bayesian inference of *Chara braunii* g41182 beta amylase homolog phylogeny. Selected homologous proteins **(Supplementary Table S1)** were aligned via M-Coffee and analyzed using BEAST 2. The depicted maximum clade credibility tree, generated via tree prior yule model and 1e4 logged MCMC chain length of 1e6, was built using TreeAnnotator with 50% burnin and a posterior probability limit of 0.5 for median node heights. The generated dendrogram was visualised using FigTree with posterior probabilities labelled on their respective nodes. The position of the putative *Chara braunii* g41182 beta amylase homolog is boxed. A protein from the cyanobacterium *Leptolyngbya* served as outgroup.

The most-closely related proteins to the protein encoded by g41182 is, again, a homolog from *K. nitens* (52/69% identical/similar amino acids in a 485 residues-long overlap with protein GAQ79925) and homologs from *Chlorella vulgaris, Chlamydomonas reinhardtii* and *Closterium* sp. NIES-68 (**Figure 6**). However, these form a shallow sister group to proteins from land plants, indicating a closer relationship to homologs in land plants (for example, (49/61% identical/similar amino acids in a 505 residues-long overlap between the *Chara braunii* g41182 gene product and protein KAJ0630873 from sunflower (*Helianthus annuus*).

## CONCLUSIONS

Here we provide insight into the enriched transcriptome of *Chara* central and nodal cells under optimised, basal growth conditions. The two transcriptome samples differ enormously from each other. While the total thallus sample showed high expression of genes associated with photosynthesis, chloroplasts and primary metabolism, the analysis of the nodal and central cell sample indicated the higher expression of genes associated with signalling, protein biosynthesis, starch metabolism and ethylene signalling pathways.

## DATA AVAILABILITY

Raw Illumina RNA-seq data produced in this study have been deposited in the DDBJ Sequence Read Archive (DRA) with the accession numbers SAMN32760189 and SAMN32760190 under BioProject PRJNA924555).

## AUTHOR CONTRIBUTIONS

WRH conceptualized the study. AH established the dissecting method, DH and AH dissected nodal and central cells, DH generated RNA samples and performed all bioinformatic analyses. DH and WRH drafted and AH edited the manuscript and all authors read and approved the final manuscript.

## FUNDING

This work was supported by the Deutsche Forschungsgemeinschaft (DFG) priority program 2237 “MAdLand”, http://madland.science, grant HE2544/18-1 to WRH and by the Open Access Publication Fund of the University of Freiburg.

The Galaxy server that was used for some calculations is in part funded by the Collaborative Research Centre 992 Medical Epigenetics (DFG grant SFB 992/1 2012) and the German Federal Ministry of Education and Research (BMBF grants 031 A538A/A538C RBC, 031L0101B/031L0101C de.NBI-epi and 031L0106 de.STAIR (de.NBI)).

## ACKNOWLEDGMENTS

We thank the Freiburg Galaxy team for running this great infrastructure, Madeleine Tarika Maier for maintaining vegetative cultures of *Chara braunii*, Manuel Brenes-Álvarez for assistance with bioinformatic analyses and the research group of Annegret Wilde (all University of Freiburg) for use of their mixer mill.

## Notes

### Competing Interest Statement

The authors have declared no competing interest.

## REFERENCES

Andrews, S. (2010). FastQC: a quality control tool for high throughput sequence data. Available at: https://www.bioinformatics.babraham.ac.uk/projects/fastqc/ [Accessed January 13, 2023].

Arribat, Y., Grepper, D., Lagarrigue, S., Qi, T., Cohen, S., and Amati, F. (2020). Spastin mutations impair coordination between lipid droplet dispersion and reticulum. PLOS Genetics 16, e1008665. doi: 10.1371/journal.pgen.1008665.

Batut, B., Freeberg, M., Heydarian, M., Erxleben, A., Videm, P., Blank, C., et al. (2022). Reference-based RNA-Seq data analysis. Galaxy Training Network. Available at: https://training.galaxyproject.org/training-material/topics/transcriptomics/tutorials/ref-based/tutorial.html [Accessed January 13, 2023].

Batut, B., Hiltemann, S., Bagnacani, A., Baker, D., Bhardwaj, V., Blank, C., et al. (2018). Community-driven data analysis training for biology. Cell Systems 6, 752-758.e1. doi: 10.1016/j.cels.2018.05.012.

Beilby, M. J. (2016). Multi-scale characean experimental system: from electrophysiology of membrane transporters to cell-to-cell connectivity, cytoplasmic streaming and auxin metabolism. Front. Plant Sci. 7. doi: 10.3389/fpls.2016.01052.

Bonnot, C., Hetherington, A. J., Champion, C., Breuninger, H., Kelly, S., and Dolan, L. (2019). Neofunctionalisation of basic helix−loop−helix proteins occurred when embryophytes colonised the land. New Phytologist 223, 993–1008. doi: 10.1111/nph.15829.

Bouckaert, R., Vaughan, T. G., Barido-Sottani, J., Duchêne, S., Fourment, M., Gavryushkina, A., et al. (2019). BEAST 2.5: An advanced software platform for Bayesian evolutionary analysis. PLOS Computational Biology 15, e1006650. doi: 10.1371/journal.pcbi.1006650.

Braun, M., and Limbach, C. (2006). Rhizoids and protonemata of characean algae: model cells for research on polarized growth and plant gravity sensing. Protoplasma 229, 133–142. doi: 10.1007/s00709-006-0208-9.

Bréhélin, C., Meyer, E. H., de Souris, J.-P., Bonnard, G., and Meyer, Y. (2003). Resemblance and dissemblance of Arabidopsis type II peroxiredoxins: similar sequences for divergent gene expression, protein localization, and activity. Plant Physiology 132, 2045–2057. doi: 10.1104/pp.103.022533.

Buschmann, H. (2020). Into another dimension: how streptophyte algae gained morphological complexity. Journal of Experimental Botany 71, 3279–3286. doi: 10.1093/jxb/eraa181.

Casanova, M. T. (2015). A revision of Chara sect. Charopsis (Characeae: Charophyceae) in Australia, including specimens collected for Bush Blitz. Aust. Systematic Bot. 27, 403–414. doi: 10.1071/SB14032.

Chao, Q., Rothenberg, M., Solano, R., Roman, G., Terzaghi, W., and Ecker†, J. R. (1997). Activation of the ethylene gas response pathway in Arabidopsis by the nuclear protein ETHYLENE-INSENSITIVE3 and related proteins. Cell 89, 1133–1144. doi: 10.1016/S0092-8674(00)80300-1.

Cheng, S., Xian, W., Fu, Y., Marin, B., Keller, J., Wu, T., et al. (2019). Genomes of subaerial Zygnematophyceae provide insights into land plant evolution. Cell 179, 1057–1067.e14. doi: 10.1016/j.cell.2019.10.019.

Chomczynski, P., and Sacchi, N. (1987). Single-step method of RNA isolation by acid guanidinium thiocyanate-phenol-chloroform extraction. Analytical Biochemistry 162, 156–159. doi: 10.1016/0003-2697(87)90021-2.

Chomczynski, P., and Sacchi, N. (2006). The single-step method of RNA isolation by acid guanidinium thiocyanate–phenol–chloroform extraction: twenty-something years on. Nat Protoc 1, 581–585. doi: 10.1038/nprot.2006.83.

Di Tommaso, P., Moretti, S., Xenarios, I., Orobitg, M., Montanyola, A., Chang, J.-M., et al. (2011). T-Coffee: a web server for the multiple sequence alignment of protein and RNA sequences using structural information and homology extension. Nucleic Acids Res 39, W13–17. doi: 10.1093/nar/gkr245.

Diao, Y., Guo, X., Li, Y., Sun, K., Lu, L., Jiang, L., et al. (2012). Pax3/7BP is a Pax7-and Pax3-binding protein that regulates the proliferation of muscle precursor cells by an epigenetic mechanism. Cell Stem Cell 11, 231–241. doi: 10.1016/j.stem.2012.05.022.

Dobin, A., Davis, C. A., Schlesinger, F., Drenkow, J., Zaleski, C., Jha, S., et al. (2013). STAR: ultrafast universal RNA-seq aligner. Bioinformatics 29, 15–21. doi: 10.1093/bioinformatics/bts635.

Domozych, D. S., and Bagdan, K. (2022). The cell biology of charophytes: Exploring the past and models for the future. Plant Physiology 190, 1588–1608. doi: 10.1093/plphys/kiac390.

Ewels, P., Magnusson, M., Lundin, S., and Käller, M. (2016). MultiQC: summarize analysis results for multiple tools and samples in a single report. Bioinformatics 32, 3047–3048. doi: 10.1093/bioinformatics/btw354.

Ferreyra, M. L. F., Pezza, A., Biarc, J., Burlingame, A. L., and Casati, P. (2010). Plant L10 Ribosomal Proteins Have Different Roles during Development and Translation under Ultraviolet-B Stress. Plant Physiology 153, 1878–1894. doi: 10.1104/pp.110.157057.

Fürst-Jansen, J. M. R., de Vries, S., and de Vries, J. (2020). Evo-physio: on stress responses and the earliest land plants. Journal of Experimental Botany 71, 3254–3269. doi: 10.1093/jxb/eraa007.

García, A. (1994). Charophyta: their use in paleolimnology. J Paleolimnol 10, 43–52. doi: 10.1007/BF00683145.

Garcia, M., Myouga, F., Takechi, K., Sato, H., Nabeshima, K., Nagata, N., et al. (2008). An Arabidopsis homolog of the bacterial peptidoglycan synthesis enzyme MurE has an essential role in chloroplast development. Plant J 53, 924–934. doi: 10.1111/j.1365-313X.2007.03379.x.

Gmelin, K. C. (1826). Flora Badensis alsatica: et confinium regionum Cis et Transrhenana plantas a Lacu Bodamico usque ad confluentem Mosellae et Rheni sponte nascentes exhibens secundum systema sexuale cum iconibus ad naturam delineatis. Mülleriana.

Hamada, T., Ueda, H., Kawase, T., and Hara-Nishimura, I. (2014). Microtubules contribute to tubule elongation and anchoring of endoplasmic reticulum, resulting in high network complexity in Arabidopsis. Plant Physiology 166, 1869–1876. doi: 10.1104/pp.114.252320.

Holzhausen, A., Stingl, N., Rieth, S., Kühn, C., Schubert, H., and Rensing, S. A. (2022). Establishment and optimization of a new model organism to study early land plant evolution: Germination, cultivation and oospore variation of Chara braunii Gmelin, 1826. Frontiers in Plant Science 13. doi: 10.3389/fpls.2022.987741.

Hopkins, B. L., and Neumann, C. A. (2019). Redoxins as gatekeepers of the transcriptional oxidative stress response. Redox Biol 21, 101104. doi: 10.1016/j.redox.2019.101104.

Kawai, H., Hanyuda, T., Akita, S., and Uwai, S. (2022). The macroalgal culture collection in Kobe University (KU-MACC) and a comprehensive molecular phylogeny of macroalgae based on the culture strains. Applied Phycology 3, 159–166. doi: 10.1080/26388081.2020.1745685.

Kuczewski, O. (1906). Morphologische und biologische Untersuchungen an Chara delicatu f. bulbillifera A. Braun. Arbeiten aus dem Laboratorium für allgemeine Botanik und Pflanzenphysiologie der Universität Zürich, 1–51.

Lan, T., Xiong, W., Chen, X., Mo, B., and Tang, G. (2022). Plant cytoplasmic ribosomal proteins: an update on classification, nomenclature, evolution and resources. The Plant Journal 110, 292–318. doi: 10.1111/tpj.15667.

Liu, R., Xu, Y.-H., Jiang, S.-C., Lu, K., Lu, Y.-F., Feng, X.-J., et al. (2013). Light-harvesting chlorophyll a/b-binding proteins, positively involved in abscisic acid signalling, require a transcription repressor, WRKY40, to balance their function. J Exp Bot 64, 5443–5456. doi: 10.1093/jxb/ert307.

Lopez, C. D., Martinovsky, G., and Naumovski, L. (2002). Inhibition of cell death by ribosomal protein L35a. Cancer Letters 180, 195–202. doi: 10.1016/S0304-3835(02)00024-1.

Lu, B., Xu, C., Awai, K., Jones, A. D., and Benning, C. (2007). A Small ATPase Protein of Arabidopsis, TGD3, Involved in Chloroplast Lipid Import *. Journal of Biological Chemistry 282, 35945–35953. doi: 10.1074/jbc.M704063200.

Martin, M. (2011). Cutadapt removes adapter sequences from high-throughput sequencing reads. EMBnet.journal 17, 10–12. doi: 10.14806/ej.17.1.200.

Martin, W. F., and Allen, J. F. (2018). An algal greening of land. Cell 174, 256–258. doi: 10.1016/j.cell.2018.06.034.

Moin, M., Saha, A., Bakshi, A., Madhav, M. S., and Kirti, P. (2021). Constitutive expression of Ribosomal Protein L6 modulates salt tolerance in rice transgenic plants. Gene 789, 145670. doi: 10.1016/j.gene.2021.145670.

Moody, L. A. (2020). Three-dimensional growth: a developmental innovation that facilitated plant terrestrialization. J Plant Res 133, 283–290. doi: 10.1007/s10265-020-01173-4.

Nishiyama, T., Sakayama, H., de Vries, J., Buschmann, H., Saint-Marcoux, D., Ullrich, K. K., et al. (2018). The Chara genome: secondary complexity and implications for plant terrestrialization. Cell 174, 448–464.e24. doi: 10.1016/j.cell.2018.06.033.

Notredame, C., Higgins, D. G., and Heringa, J. (2000). T-Coffee: A novel method for fast and accurate multiple sequence alignment. J Mol Biol 302, 205–217. doi: 10.1006/jmbi.2000.4042.

Paysan-Lafosse, T., Blum, M., Chuguransky, S., Grego, T., Pinto, B. L., Salazar, G. A., et al. (2023). InterPro in 2022. Nucleic Acids Research 51, D418–D427. doi: 10.1093/nar/gkac993.

Pfalz, J., and Pfannschmidt, T. (2013). Essential nucleoid proteins in early chloroplast development. Trends in Plant Science 18, 186–194. doi: 10.1016/j.tplants.2012.11.003.

Pinto, F. L., Thapper, A., Sontheim, W., and Lindblad, P. (2009). Analysis of current and alternative phenol based RNA extraction methodologies for cyanobacteria. BMC Molecular Biology 10, 79. doi: 10.1186/1471-2199-10-79.

Rambaut, A. (2018). FigTree software, version 1.4.4. Available at: http://tree.bio.ed.ac.uk/software/figtree/ [Accessed February 10, 2023].

Rambaut, A., Drummond, A. J., Xie, D., Baele, G., and Suchard, M. A. (2018). Posterior summarization in bayesian phylogenetics using tracer 1.7. Systematic Biology 67, 901–904. doi: 10.1093/sysbio/syy032.

Ramu, V. S., Dawane, A., Lee, S., Oh, S., Lee, H.-K., Sun, L., et al. (2020). Ribosomal protein QM/RPL10 positively regulates defence and protein translation mechanisms during nonhost disease resistance. Molecular Plant Pathology 21, 1481–1494. doi: 10.1111/mpp.12991.

Rekha, A., and Sujathamma, P. (2020). Biochemical, phytochemical and antibacterial analysis of Chara braunii c.c. gmelin. International Journal of Botany Studies 5, 358–360.

Ruhfel, B. R., Gitzendanner, M. A., Soltis, P. S., Soltis, D. E., and Burleigh, J. G. (2014). From algae to angiosperms–inferring the phylogeny of green plants (Viridiplantae) from 360 plastid genomes. BMC Evolutionary Biology 14, 23. doi: 10.1186/1471-2148-14-23.

Sato, M., Sakayama, H., Sato, M., Ito, M., and Sekimoto, H. (2014). Characterization of sexual reproductive processes in Chara braunii (Charales, Charophyceae). Phycological Research 62, 214–221. doi: 10.1111/pre.12056.

Scholz, I., Lott, S. C., Behler, J., Gärtner, K., Hagemann, M., and Hess, W. R. (2019). Divergent methylation of CRISPR repeats and cas genes in a subtype I-D CRISPR-Cas-system. BMC Microbiology 19, 147.1-147.11.

Schubert, H., Holzhausen, A., and Nowak, P. (2016). “Individualentwicklung der Characeen,” in Armleuchteralgen: Die Characeen Deutschlands (Berlin, Heidelberg: Springer), 57–78. doi: 10.1007/978-3-662-47797-7_6.

Solano, R., Stepanova, A., Chao, Q., and Ecker, J. R. (1998). Nuclear events in ethylene signaling: a transcriptional cascade mediated by ETHYLENE-INSENSITIVE3 and ETHYLENE-RESPONSE-FACTOR1. Genes Dev. 12, 3703–3714. doi: 10.1101/gad.12.23.3703.

Tagami, S., Ohnishi, K., Hikichi, Y., and Kiba, A. (2021). Trigalactosyldiacylglycerol 3 protein orthologs are required for basal disease resistance in Nicotiana benthamiana. Plant Biotechnol (Tokyo) 38, 373–378. doi: 10.5511/plantbiotechnology.21.0624a.

Tang, D., Christiansen, K. M., and Innes, R. W. (2005). Regulation of plant disease resistance, stress responses, cell death, and ethylene signaling in Arabidopsis by the EDR1 protein kinase. Plant Physiol 138, 1018–1026. doi: 10.1104/pp.105.060400.

The Galaxy Community (2022). The Galaxy platform for accessible, reproducible and collaborative biomedical analyses: 2022 update. Nucleic Acids Research 50, W345–W351. doi: 10.1093/nar/gkac247.

Vries, J. de, and Archibald, J. M. (2018). Plant evolution: landmarks on the path to terrestrial life. New Phytologist 217, 1428–1434. doi: 10.1111/nph.14975.

Wawrzynska, A., Christiansen, K. M., Lan, Y., Rodibaugh, N. L., and Innes, R. W. (2008). Powdery mildew resistance conferred by loss of the ENHANCED DISEASE RESISTANCE1 protein kinase Is suppressed by a missense mutation in KEEP ON GOING, a regulator of abscisic acid signaling. Plant Physiology 148, 1510–1522. doi: 10.1104/pp.108.127605.

Wickett, N. J., Mirarab, S., Nguyen, N., Warnow, T., Carpenter, E., Matasci, N., et al. (2014). Phylotranscriptomic analysis of the origin and early diversification of land plants. PNAS 111, E4859–E4868. doi: 10.1073/pnas.1323926111.

Xiong, W., Lan, T., and Mo, B. (2021). Extraribosomal Functions of Cytosolic Ribosomal Proteins in Plants. Frontiers in Plant Science 12. Available at: https://www.frontiersin.org/articles/10.3389/fpls.2021.607157 [Accessed February 11, 2023].

Young, M. D., Wakefield, M. J., Smyth, G. K., and Oshlack, A. (2010). Gene ontology analysis for RNA-seq: accounting for selection bias. Genome Biology 11, R14. doi: 10.1186/gb-2010-11-2-r14.

Zhang, Z., Ma, X., Liu, Y., Yang, L., Shi, X., Wang, H., et al. (2022). Origin and evolution of green plants in the light of key evolutionary events. Journal of Integrative Plant Biology 64, 516–535. doi: 10.1111/jipb.13224.

